# Leaf variegation in a barley EMS mutant is controlled by two epistatic mutations

**DOI:** 10.64898/2025.12.15.694392

**Authors:** Hélène Pidon, Shobhashree Rajulareddy Nagireddy, Michael Melzer, Axel Himmelbach, Nils Stein

## Abstract

Variegation mutants provide valuable insights into chloroplast biogenesis. We characterized a newly identified variegated barley mutant, in which the phenotype is controlled by duplicate dominant epistasis—representing, to our knowledge, the first reported case of digenic control in chloroplast-deficient mutants. The causal loci, *Var4* and *Var5*, were mapped on chromosomes 2H and 3H. We used whole-genome resequencing to identify candidate genes. Our two top candidate genes are an *NBR1-like selective autophagy receptor* gene and a *DNAJ-domain containing* gene, respectively. Based on their homology-based functional annotation, both candidates could be implicated in chloroplast proteostasis, regulating protein import, folding, and/or degradation. We propose that mild, independent defects in proteostasis from each mutation act synergistically to surpass a functional threshold, impairing chloroplast development in early leaves while allowing partial recovery in later stages. These findings highlight a novel digenic mechanism underlying variegation and point to proteostasis as a central vulnerability in chloroplast biogenesis.

**Highlight:** We identified digenic control of leaf variegation in a barley mutant. Candidate genes, identified through whole-genome resequencing, suggest roles in chloroplast proteostasis.

## Introduction

Since the 1950s, the induction of mutations in plant populations has been extensively used, either to obtain new traits or to study the genetic basis of a phenotypic defect to shed light on developmental processes. In this frame, the study of photosynthesis-related mutants exhibiting abnormal leaf coloration is of particular interest to study chloroplast development (Taylor *et al*., 1987; Barkan, 1998; Leon *et al*., 1998).

Chloroplast biogenesis is the process by which chloroplasts differentiate from proplastids (Pogson and Albrecht, 2011). This complex process relies on signaling between the nucleus and the chloroplasts and requires both plastid and nuclear genes. Out of approximately 3,000 chloroplast proteins, less than 100 are encoded by the plastome (Li and Chiu, 2010; Petersen *et al*., 2013). Most mutations resulting in aberrant leaf coloration are therefore affecting nuclear genes, and their identification and functional characterization brings us one important step closer to the understanding of regulatory mechanisms of chloroplast biogenesis.

Among phenotypes with chlorophyll deficiency, leaf variegation is particularly interesting. It is characterized by areas of the leaf with a certain level of pigment deficiency, either in patches in dicotyledons or stripes in monocotyledons (Yu *et al*., 2007). Variegated mutants usually have well-developed chloroplasts in green leaf sectors and defective ones in the pigment-deficient sectors. An important advantage of variegated mutants over albinos (complete chlorophyll-deficiency) is that they usually survive and produce seeds, which enables genetic studies. Somatic chimerism can be the reason for leaf variegation, where cells of each sector present a different genotype. However, in the case of mutants, the genotype is most likely identical in the whole plant.

Several variegation genes have been cloned and characterized through the years, both in *Arabidopsis thaliana* (Wu *et al*., 1999; Chen *et al*., 2000; Takechi *et al*., 2000; Sakamoto *et al*., 2002; Wang *et al*., 2004; Zheng *et al*., 2016; Zagari *et al*., 2017), and in other plant species like rice (Hayashi-Tsugane *et al*., 2014; Li *et al*., 2018), barley (Li *et al*., 2019; Overlander-Chen *et al*., 2024), or tomato (Song *et al*., 2023; Dechkrong *et al*., 2024). The best-described genes are *Arabidopsis thaliana VAR1* and *VAR2*, encoding genes of the filamentation temperature-sensitive (FtsH) metalloprotease family (Lindahl *et al*., 1996, 2000; Chen *et al*., 2000; Takechi *et al*., 2000; Sakamoto *et al*., 2002). The functional model for these genes describes two pairs of FtsH proteins, FtsH5/VAR1 and FtsH2/VAR2, that form oligomeric complexes (Yu *et al*., 2004). In the case of *VAR2*, a threshold model explains variegation, FtsH8 compensating for the lack of functional VAR2 in *var2* mutants. In barley, three variegation mutations were studied in greater detail: *albostrians*, caused by a mutation in *Hordeum vulgare CCT Motif Family gene 7* (*HvCMF7*) (Li *et al*., 2019), *luteostrians*, for which the *ATP-Dependent Clp Protease Subunit C1* has been postulated as a convincing candidate gene (Li *et al*., 2021), and *grandpa1.a*, caused by a large deletion in a gene coding for plastid terminal oxidase homologous to *A. thaliana IMMUTANS* (Overlander-Chen *et al*., 2024). More barley variegated mutants are listed in the NordGen database (Hansson *et al*., 2024; https://bgs.nordgen.org/), such as *var1* to *var3* (variegated 1 to 3), *wst1* to *wst7* (White streak 1 to 7), and *yst1* to *yst5* (Yellow streak 1 to 5).

Here we studied the genetic basis of a leaf variegation phenotype induced by the alkylating agent ethyl-methane sulfonate (EMS) in the genetic background of the two-rowed winter barley cultivar ‘Igri’. We showed that, unlike previously identified variegation traits, variegation is here controlled by two epistatic genes, which we named Variegated 4 (*Var4*) and Variegated 5 (*Var5*), respectively, in accordance with the naming conventions of barley gene (Hansson *et al*., 2024). Performing whole-genome sequencing of variegated and green progenies of the mutant x wild type backcross, we narrowed down a small number of candidate genes for the two loci, which hint to chloroplast proteostasis as the developmental process that is centrally affected by the mutation.

## Materials and methods

### Plant material

The variegated parent is a fourth-generation selfed (M4) mutant from an EMS-mutagenized population of the barley cultivar ‘Igri’, obtained according to the protocol of Gottwald et al. (2009) with 40 mM of EMS treatment for 16 hours. Three F_2_ populations were constructed: the cultivar ‘Alraune’ was used as ‘wild-type’ (WT) in reciprocal crosses to establish the populations ‘AM’ (Alraune x mutant) and ‘MA’ (mutant x Alraune), respectively. Reciprocal crosses were initiated to test for nucleus- or cytoplasm-controlled variegation. In addition, the true WT cultivar ‘Igri’ was used for backcrossing to form the ‘IM’ (Igri x mutant) population. F_3_ progenies of four and two variegated F_2_ from the AM and MA populations, respectively, were grown to confirm the segregation.

All plants were cultivated in a greenhouse under a day/night temperature cycle of 20°C/15°C and a photoperiod of 16 hours of light and 8 hours of darkness. Natural light was extended when necessary using incandescent lamps (SON-T Agro 400, MASSIVEGROW) at 300 μmol photons m^−2^ s^−1^.

### Phenotyping and Ultrastructural Analysis

The presence of stripes on the leaves was assessed as a qualitative trait 10 days to three months after sowing , such that a single variegated leaf was sufficient to classify the plant as variegated. The extent of variegation was not assessed by any quantitative measurement.

Leaves of the mutant and WT Igri plants were collected 14 days and 6 months after sowing and prepared for morphological and ultrastructural analysis. For this purpose, cuttings of 1-2 mm² of the central part of the leaves of at least 5 plants were fixed with glutaraldehyde and paraformaldehyde, embedded in resin, sectioned, and examined by light and transmission electron microscopy (TEM) according to the protocol of (Li *et al*., 2021).

### Genotyping-by-sequencing (GBS)

To perform linkage mapping, 269 and 183 F_3_ plants from AM and MA populations, respectively, were genotyped using genotyping-by-sequencing. Genomic DNA was extracted according to the guanidine isothiocyanate-based protocol described by Milner et al. (2019). GBS library preparation using *PstI* and *MspI* enzymes and sequencing followed essentially a previously described procedure (Zhang *et al*., 2024). Sequencing (single-read, SP flowcell with 100-cycle kit) was performed using a custom sequencing primer (Wendler et al., 2014) and the Illumina NovaSeq6000 device (Illumina, Inc., San Diego, CA, USA) at IPK-Gatersleben.

The GBS sequencing data were aligned to ‘Morex’ reference genome version 3 (Mascher *et al*., 2021), and variants were called as described by Milner et al. (2019). The variants table was filtered as followed: homozygous and heterozygous variant calls with a depth of sequencing inferior to 4 and 6, respectively, were filtered out, variant sites were accepted wherever they had a minimum mapping quality score of 40, a maximum fraction of 20% of missing data, a maximum of 60% of heterozygous call, and a minimum allele frequency of 30%. GBS data from the parents was used to assign each call to its respective parental allele.

### Linkage mapping

The genotype matrix obtained from GBS was additionally filtered to construct the linkage map. Only variant sites with less than 15% missing data and individuals with less than 25% missing data were retained. The data were manually curated to remove suspicious double crossing overs.

The linkage map was built in R 4.1.3 with the package ASmap (Taylor and Butler, 2017), using the mstmap.cross function with the chromosome information from the GBS mapping as linkage group and a high p-value of 1e-2 to avoid splitting them into several linkage groups, as well as the option “anchor=TRUE” to respect the order of the markers on the chromosomes. The linkage map was visualized with the package LinkageMapView (Ouellette *et al*., 2018) and edited in Inkscape 1.1.2. Finally, the linkage mapping of the variegation trait was performed with the R/qtl package (Broman *et al*., 2003) using single maximum likelihood-based interval mapping, performed via the EM algorithm and with a binary phenotype model, as well as composite interval mapping. The significance threshold was calculated based on 1000 permutations at a 5% alpha level, and the interval was determined based on 1-LOD drop.

### PACE genotyping

For a finer delimitation of the mapping intervals, polymorphisms identified in the candidate intervals in the GBS data from the parental lines, Alraune and the mutant, were developed into PCR Allele Competitive Extension (PACE) markers through 3CR Bioscience (Essex, UK) free assay design service (Supplementary Table S1). Genotyping with the PACE markers was carried out as described by Pidon et al. (2020) on 89 selected F_2_ plants, of which 21 were variegated, from AM and MA populations, and the parents of the populations.

### Whole-Genome Shotgun Sequencing and polymorphisms analysis

A total of 30 F2 individuals from the IM population, of which six exhibited a striped phenotype, were selected for whole-genome shotgun sequencing (WGS). DNA was extracted from young leaf tissue using the Milner et al. (2019) protocol. Libraries were prepared with Illumina Nextera DNA Flex Kit following the standard protocols from the manufacturer (Illumina, Inc., San Diego, CA, USA), quantified by qPCR (Mascher *et al*., 2013), and sequenced on the NovaSeq6000 platform using the XP workflow (S1 flowcell, paired-end, 2 x 151 cycles). High-quality reads were then mapped against two barley genomes: the reference genome Morex (Mascher *et al*., 2021) and the WT parent Igri (Jayakodi *et al*., 2024) as described above for GBS data.

Intervals of interest on the Igri genome were identified by blasting the flanking 100 bp of each Morex interval. SNPs located on chromosomal regions of interest were extracted using VCFtools (Danecek *et al*., 2011). Annotation of polymorphisms was carried out using SnpEff (Cingolani *et al*., 2012), employing the respective genome annotations. The variants were then filtered as follows: : only SNPs with a single alternate allele were retained; among the six striped individuals, none could be fixed for the reference allele, and they had to include no more than one heterozygous genotype and at least two homozygous alternate genotypes; among the 24 non-striped individuals, at least one had to be homozygous for the reference allele and at least one homozygous for the alternate allele. Genes were defined as candidates if they displayed a filtered SNPs with ‘MODERATE’ or ‘HIGH’ predicted effects on protein , and were not classified as low confidence (LC) in the Morex genome, annotated as transposable elements, or had significant similarity to transposons based on BLAST searches.

The homologous genes in *Arabidopsis thaliana* were retrieved based on the TAIR annotation displayed in Panbarlex (https://panbarlex.ipk-gatersleben.de/; Jayakodi et al., 2024). Orthologs in other monocotyledonous species were retrieved by identifying the orthogroups to which the genes belong in Monocots PLAZA 5.0 (Van Bel *et al*., 2022). The orthologs were then aligned, and the conservation of the amino acid mutated in the EMS mutant was assessed. The sequence logos in the 101 bp interval around these polymorphisms were generated using WebLogo 3 (Crooks *et al*., 2004).

## Results

### Description and segregation of the variegated phenotype

The mutant plants display a white-striped variegated phenotype with different levels of penetration, from a mostly white plant to a low number of narrow white stripes (Fig. 1). As the plant grows, tillers present a different level of variegation. After 6 months of cultivation, the stripes are progressively turning yellow (Fig. 2). The F_3_ progenies of variegated F_2_ plants showed a similar but not as pronounced phenotype as the most variegated mutant plants. Stripes sometimes occurred only on secondary tillers.

**Fig. 1:**
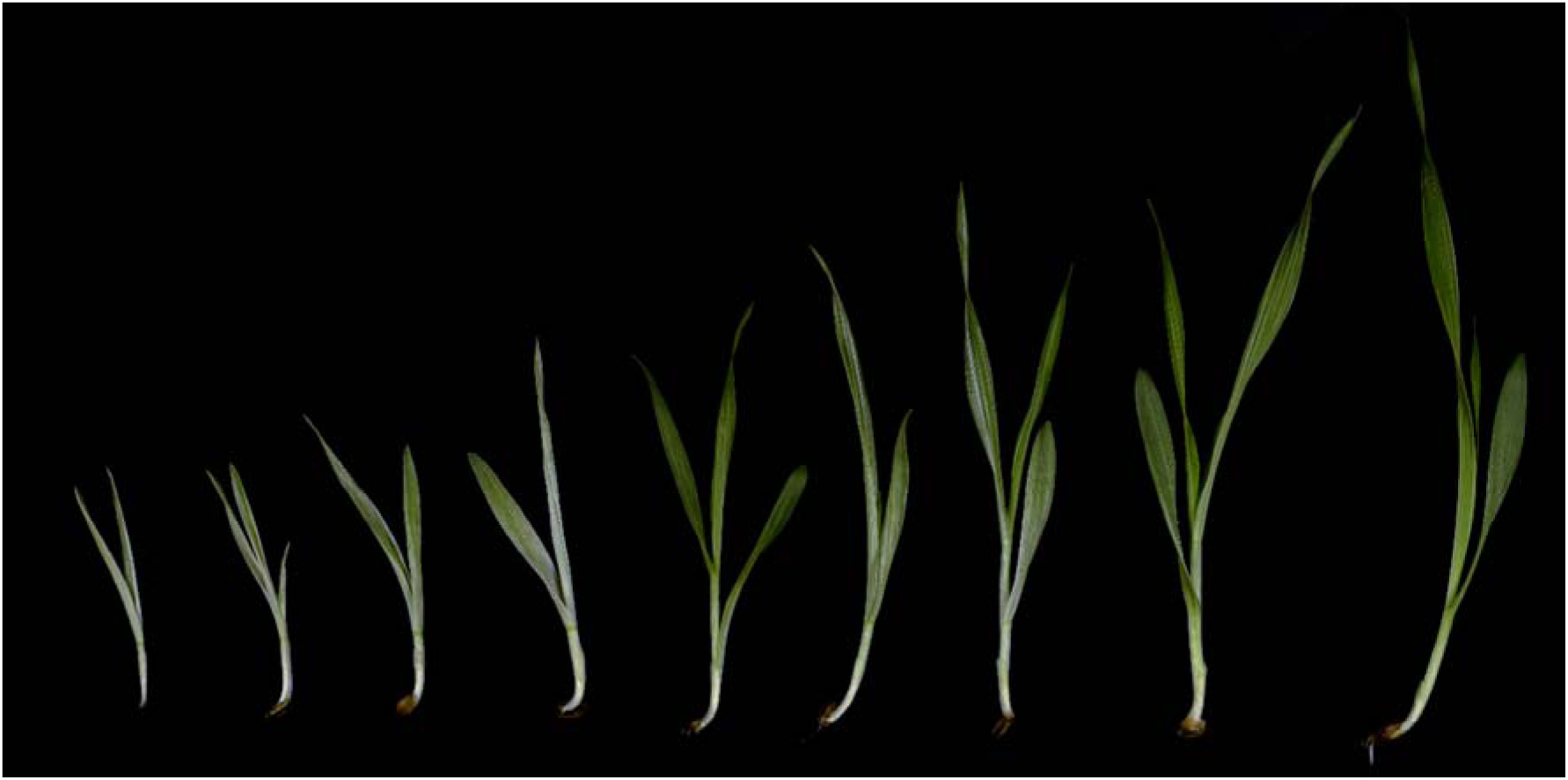
Penetration of the mutant phenotype on 14-day-old mutant plants. The mutant plantlets displayed diverse levels of variegation, from almost albino plantlets to plantlets with only a few white stripes. The level of penetration of the phenotype also affects growth.

**Fig. 2:**
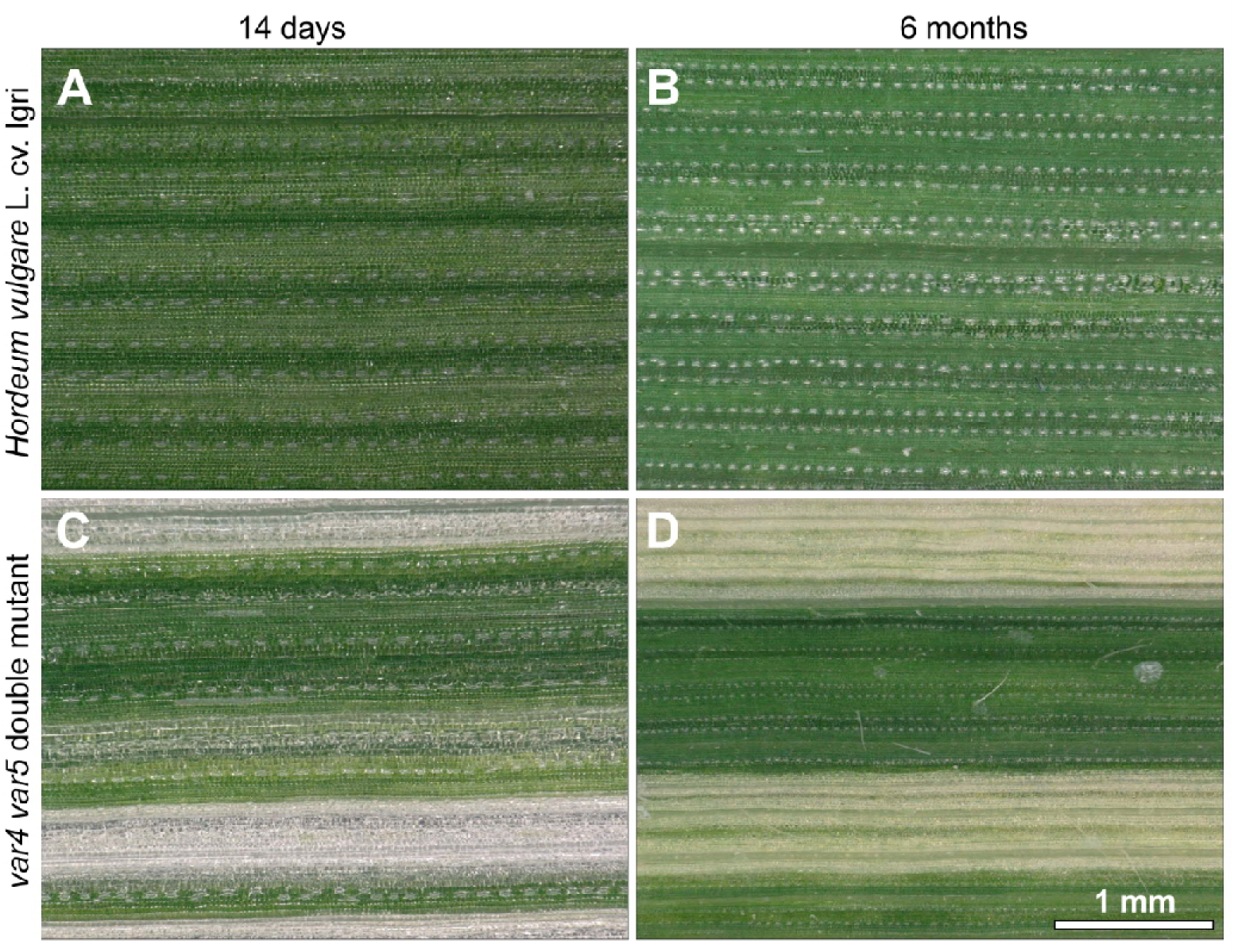
Digital microscopy observation of the central part of leaves from WT and mutant Igri plants 14 days (A, C) and 6 months (B, D) after sowing. Stripes are composed of cells from an identical cell line. After 6 months, the stripes are turning yellow, suggesting a partial recovery.

To assess if there is a maternal inheritance of the variegation, the proportion of variegated plants in the reciprocal crosses AM and MA was compared. These proportions were 17 out of 298 and 6 out of 203 (Table 1, Supplementary Table S2), respectively, and thus are not statistically different (Chi-Square Test of Independence, χ²(1, N=501)=2.083, *p*=.149), indicating that the trait is nuclear-encoded and -inherited. The two populations are therefore treated as one for the rest of the study.

**Table 1:** Plant phenotypes in the three F_2_ populations. The number of variegated plants and the total number of plants are indicated. The Chi-square test of goodness of fit for the 1:15 ratio and the corresponding p-value in each population have been calculated.

| Population | variegated | total | $\chi^2$ goodness of fit<br>1:15 | p-value |
| --- | --- | --- | --- | --- |
| AM | 17 | 298 | 0.151 | .697 |
| MA | 6 | 203 | 3.760 | .052 |
| IM | 6 | 95 | 0.001 | .979 |

The segregation of the variegation trait in the combined AM-MA population fit the segregation ratio of 1:15, which could be expected for two recessive epistatic factors (Chi-Square Goodness of Fit Test χ²(1)=2.354, *p*=.125). The segregation in the IM population (Table 1, Supplementary Table S2) does not statistically differ from the one in the AM-MA population (Chi-Square Test of Independence, χ²(1, N=596)=0.513, *p*=.474), confirming that both genetic factors are inherited from the EMS mutant. The segregation of variegation in F_3_ was checked on the progenies of six variegated F_2_. All 12 F_3_ plants per line were variegated, confirming that the genotype is fixed at the responsible loci in variegated plants.

### Genetic mapping reveals two epistatic loci

After filtering of the GBS data, 265 and 181 plants from the AM and MA populations, respectively, and 3,022 SNPs remained (Supplementary Table S3). Simple interval mapping and composite interval mapping, performed on the combined AM and MA populations, identified two major loci, confirming the hypothesis derived from the segregation pattern (Fig. 3A, Supplementary Table S4). Those two loci, located on chromosome 2H and chromosome 3H, were named Variegated 4 (gene symbol: *Var4*) and Variegated 5 (*Var5*), respectively.

**Fig. 3:**
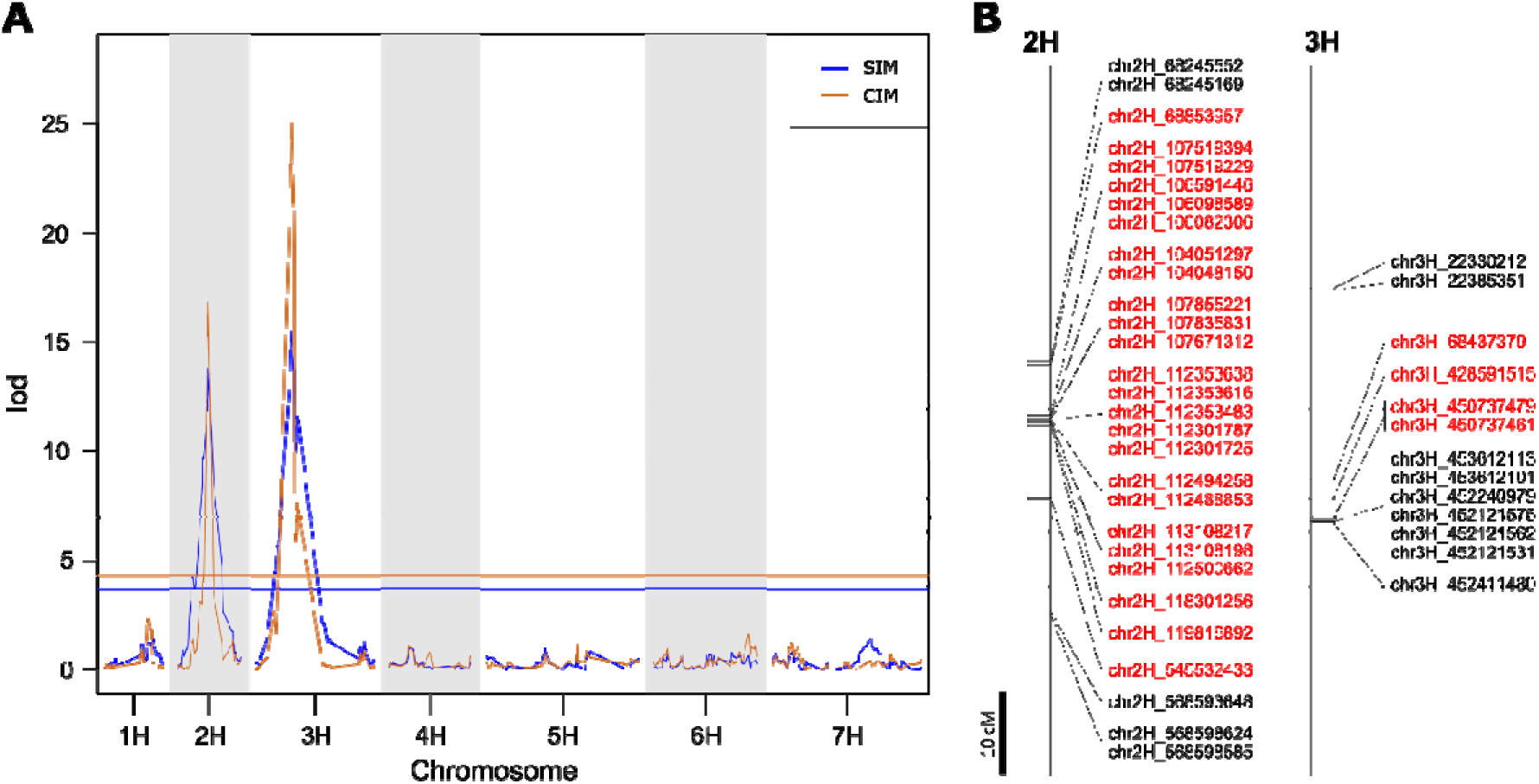
Interval mapping of the variegation trait. **A.** LOD scores calculated with single interval mapping (SIM, blue) and composite interval mapping (CIM, orange) along the chromosomes. The horizontal bars represent the 5% probability thresholds calculated based on 1000 permutations. **B.** Graphical representation of the significant intervals. The markers in red represent the intervals calculated by 1-LOD drop.

The respective recessive mutant alleles are *var4.a* and *var5.a.* Based on a 1-LOD drop, the interval of *Var4* covers 476.7 Mbp from markers chr2H_68853957 and chr2H_545532433. The interval of *Var5* spanned over 382.3 Mbp and contains markers chr3H_68437379, chr3H_428591515, chr3H_450737479, and chr3H_450737461 (Fig. 3B).

A visual inspection of the genotype and crossover matrix confirmed and refined the intervals to chr2H: bp-position 119,876,820 - 545,212,322 and chr3H: bp-position 145,112,389 - 450,737,461 intervals for *Var4* and *Var5*, respectively. Only plants homozygous for the mutant allele at both loci presented the variegated phenotype. Genotyping with PACE markers of 89 F_2_ plants confirmed those intervals, without allowing to further reduce them (Fig. 4, Supplementary Table S5).

**Fig. 4:**
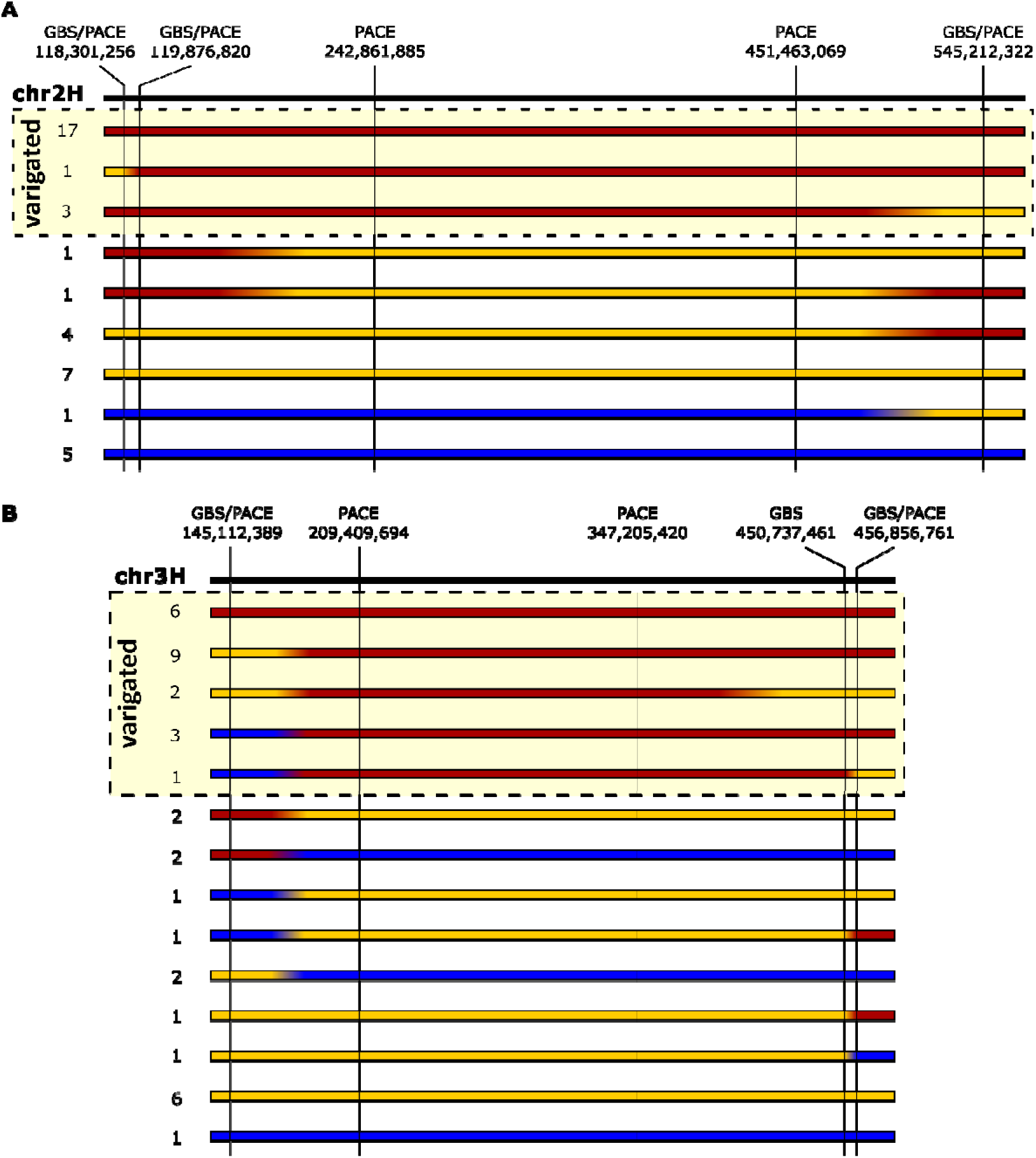
Graphical representation of the genotype at *Var4* (A) and *Var5* (B) loci of the F_2_ plants fixed for the mutant haplotype at the other locus. Vertical bars represent the position of the markers, identified by their nature (GBS, PACE, or both) and their physical position on the ‘Morex’ reference genome. Each horizontal rectangle represents a genotype in the interval. Red, blue, and yellow sections depict homozygous mutant, homozygous ‘Alraune’, and heterozygous alleles, respectively. The number on the left side of the bar shows the number of variegated F_2_ plants displaying this genotype. The plants carrying the genotypes in the dashed box displayed a variegated phenotype.

The mapping intervals of *var4* and *var5* mutations are annotated with 5,131 and 3,409 genes, including 3,909 and 2,577 protein-coding genes, of which 2,057 and 1,279 are annotated with high confidence, respectively.

### Five genes are candidates in each interval

To reduce the number of candidate genes, we performed WGS on 20 F_2_ plants from the backcross between the mutant and the WT cultivar ‘Igri’ (IM population) and compared the polymorphisms observed on the six variegated plants and 24 green ones and their effects on coding genes (Supplementary Tables S6, S7, S8, and S9). The same list of five candidate mutations on each interval was found after mapping the WGS data on Morex or Igri genomes and filtering the results for their impact on protein-coding genes annotated with high confidence (Table 2).

**Table 2:** Candidate genes and their mutation in the *var4 var5* double mutant. The coordinates of the 606 mutation are given in relation to Morex v3 Mascher *et al*., 2021) and Igri (Jayakodi *et al*., 2024) genome assemblies. The functional annotation and *A. thaliana* homolog provided are those displayed in the PanBARLEX portal (https://panbarlex.ipk-gatersleben.de/).

| Morex gene ID | Igri gene ID | Pol. coordinates<br>Morex | Pol. coordinates<br>Igri | Nucl.<br>Pol. | Typical<br>EMS? | Protein<br>Pol. | Functional annotation | *Arabidopsis*<br>*thaliana*<br>homolog |
| --- | --- | --- | --- | --- | --- | --- | --- | --- |
| **HORVU.MOREX.PROJ.2HG001239**<br>**10** | **HORVU.IGRI.PROJ.2HG001254**<br>**10** | **chr2H:20671885**<br>**8** | **chr2H:20571237**<br>**3** | **C>T** | **yes** | **A18V** | **Tetratricopeptide repeat (TPR)-like superfamily protein** | **AT5G46400 / PRP39-2** |
| HORVU.MOREX.PROJ.2HG001250<br>10 | HORVU.IGRI.PROJ.2HG001265<br>00 | chr2H:21644660<br>2 | chr2H:21541296<br>6 | T>A | no | K72* | tRNA wybutosine-synthesizing protein 2 homolog | AT4G04670 / TRM5C |
| **HORVU.MOREX.PROJ.2HG007273**<br>**00** | **HORVU.IGRI.PROJ.2HG007076**<br>**40** | **chr2H:36940587**<br>**4** | **chr2H:36626289**<br>**6** | **G>A** | **yes** | **T65I** | **Unknown protein** | **-** |
| **HORVU.MOREX.PROJ.2HG001507**<br>**00** | **HORVU.IGRI.PROJ.2HG001515**<br>**40** | **chr2H:49912441**<br>**3** | **chr2H:49423501**<br>**9** | **C>T** | **yes** | **R219K** | **Ribonuclease H-like superfamily protein** | **AT3G25270** |
| **HORVU.MOREX.PROJ.2HG001544**<br>**10** | **HORVU.IGRI.PROJ.2HG001553**<br>**10** | **chr2H:52596062**<br>**9** | **chr2H:52118707**<br>**2** | **C>T** | **yes** | **G410D** | **ZZ-type zinc finger-containing protein** | **AT4G24690 / NBR1** |
| **HORVU.MOREX.PROJ.3HG002264**<br>**20** | **HORVU.IGRI.PROJ.3HG002275**<br>**10** | **chr3H:18588293**<br>**4** | **chr3H:18702440**<br>**2** | **C>T** | **yes** | **R63C** | **phytochrome kinase substrate 4** | **AT5G04190 / PSK4** |
| **HORVU.MOREX.PROJ.3HG002388**<br>**70** | **HORVU.IGRI.PROJ.3HG002398**<br>**30** | **chr3H:34537213**<br>**3** | **chr3H:34352766**<br>**6** | **G>A** | **yes** | **S696F** | **DNAJ heat shock N-terminal domain-containing protein-like** | **AT5G18750** |
| HORVU.MOREX.PROJ.3HG002390<br>80 | HORVU.IGRI.PROJ.3HG002400<br>20 | chr3H:34735995<br>8 | chr3H:34551220<br>7 | A>T | no | K38N | F-box/FBD/LRR protein | AT1G69630 |
| **HORVU.MOREX.PROJ.3HG002446**<br>**60** | **HORVU.IGRI.PROJ.3HG002456**<br>**00** | **chr3H:40481116**<br>**7** | **chr3H:40227822**<br>**5** | **C>T** | **yes** | **T1017I** | **Histone-lysine N-methyltransferase** | **AT4G27910 / SDG16** |
| HORVU.MOREX.PROJ.3HG002467<br>00 | HORVU.IGRI.PROJ.3HG002476<br>70 | chr3H:42216747<br>0 | chr3H:41962839<br>9 | A>G | no | W119<br>R | Lectin-domain containing receptor kinase A4.3 | AT1G66235 |

Among the five candidate mutations for *var4.a*, four are typical EMS mutations (C to T or G to A) leading to amino acid substitutions, and one is a non-typical EMS mutation (T to A) causing a premature stop in a protein annotated as tRNA wybutosine-synthesizing.

Moreover, three out of five are at the heterozygous state in one of the six striped plants (MZ42_46_3_36), and there is no coverage at HORVU.MOREX.PROJ.2HG00727300 locus, annotated as an unknown protein. Only the polymorphism in HORVU.MOREX.PROJ.2HG00154410 is homozygous for the alternate allele in all six striped plants, and therefore the best candidate. This gene is one of the two closest homologs of *A. thaliana NBR1* which codes for an ubiquitin-binding autophagy receptor protein directing the autophagy of damaged chloroplasts as well as regulating chloroplast protein import through the control of translocon at the outer envelope membrane of chloroplasts (TOC) protein levels (Lee *et al*., 2023; Wan *et al*., 2023). In the *var4 var5* double mutant, the mutation replaces a glycine with an aspartic acid. This residue lies just 16 amino acids upstream of the zinc finger domain, within the linker region that positions the domain for binding. The introduction of a negatively charged residue in this position could alter the domain’s orientation or electrostatic environment, potentially reducing interaction efficiency.

All candidate mutations for *var5.a* are responsible for amino acid changes. Three of them have the typical EMS signature (HORVU.MOREX.PROJ.3HG00226420, HORVU.MOREX.PROJ.3HG00238870, and HORVU.MOREX.PROJ.3HG00244660), and are therefore better candidates than the two that do not (HORVU.MOREX.PROJ.3HG00239080 and HORVU.MOREX.PROJ.3HG00246700). HORVU.MOREX.PROJ.3HG00226420 is annotated as a phytochrome kinase substrate 4, and is homologous to *A. thaliana PKS4*, involved in phototropism and the regulation of light signaling (Schepens *et al*., 2008; Demarsy *et al*., 2012). The protein produced by HORVU.MOREX.PROJ.3HG00244660 is a histone-lysine N-methyltransferase homologous to *AtSDG16*, involved in transcriptional regulation via chromatin modification, in particular in response to drought stress (Schuettengruber *et al*., 2011; Chen *et al*., 2017; Liu *et al*., 2018). Finally, HORVU.MOREX.PROJ.3HG00238870 encodes a chaperone protein, DNAJ. DNAJ proteins are co-chaperones of HSP70s, ensuring correct folding of proteins and targeting misfolded proteins for degradation in collaboration with Clp proteases (Pulido *et al*., 2013; Pulido and Leister, 2018; Rodriguez-Concepcion *et al*., 2019). While the amino acids affected by mutations in the other two candidates are poorly conserved, the substitution in HORVU.MOREX.PROJ.3HG00238870 affects a highly conserved serine—present in 284 out of 326 proteins in the orthogroup (Supplementary Fig. S1)—replacing it with phenylalanine in one of the protein’s two DUF3444 domains . This radical change in a domain that is highly conserved across most DNAJ proteins may impair its function in the described mutant.

Beyond these polymorphisms, EMS-type mutations were identified in non-coding regions. After mapping the WGS data to the Igri genome and applying an additional filter to retain only homozygous calls in striped plants, 402 and 505 mutations were detected at the *Var4* and *Var5* loci, respectively (supplementary tables 6 to 9). Among these, 15 occur in upstream gene regions at each locus. While these may include polymorphisms in regulatory elements and cannot be ruled out as the underlying cause of the phenotype, they do not appear to impact obvious candidate genes.

### Chloroplast ultrastructure in the *var4 var5* double mutant

Chloroplasts of leaf samples of 14 days (white stripes in the mutant) and six-month-old WT and mutant plants after sowing (stripes turned yellow) were analyzed by TEM. Chloroplasts in the WT and green sectors of the mutant leaves display well-developed stroma and grana thylakoids (Fig. 5 A, B, D, E). In contrast, chloroplasts of the white sectors of 14-day-old mutant plants appear significantly smaller. In addition, remaining prolamellar bodies, poorly developed thylakoids, and a lack of grana stacks indicate impaired chloroplast biogenesis (Fig. 5C, F).

**Fig. 5:**
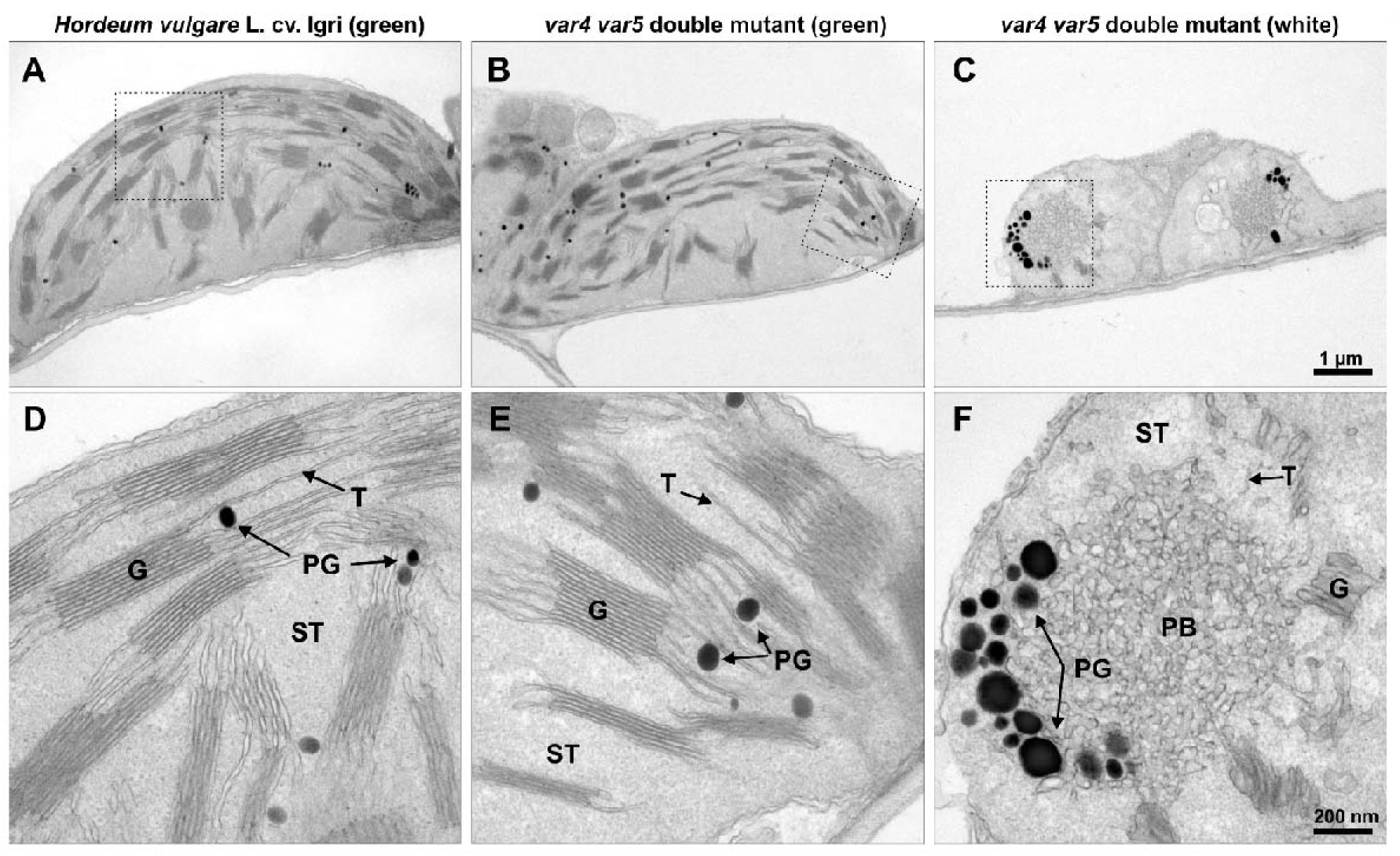
Chloroplast ultrastructure of WT Igri and *var4 var5* double mutant plants 14 days after sowing. Chloroplast with close up of WT Igri **(A, D)**, green **(B, E),** and white leaf sectors of the *var4 var5* double mutant **(C, F)**, respectively. G, grana; PB, prolamellar body, PG, plastoglobuli; ST, stroma thylakoid; T, thylakoid.

After six months of growth, chloroplasts of WT plants and green sectors of mutant variegated plants appear fully developed (Fig. 6A, B, E, F). However, in the now yellow sectors of the mutant, different levels of chloroplast development were observed, from chloroplasts with similar disorganization as observed after 14 days (Fig. 6C, G), up to chloroplasts with intermediate size and characterized by thylakoids and small grana (Fig. 6D, H). This observation suggests that chloroplast development is limited or rests at a certain level in the variegated segments of mutant leaves during plant growth.

**Fig. 6:**
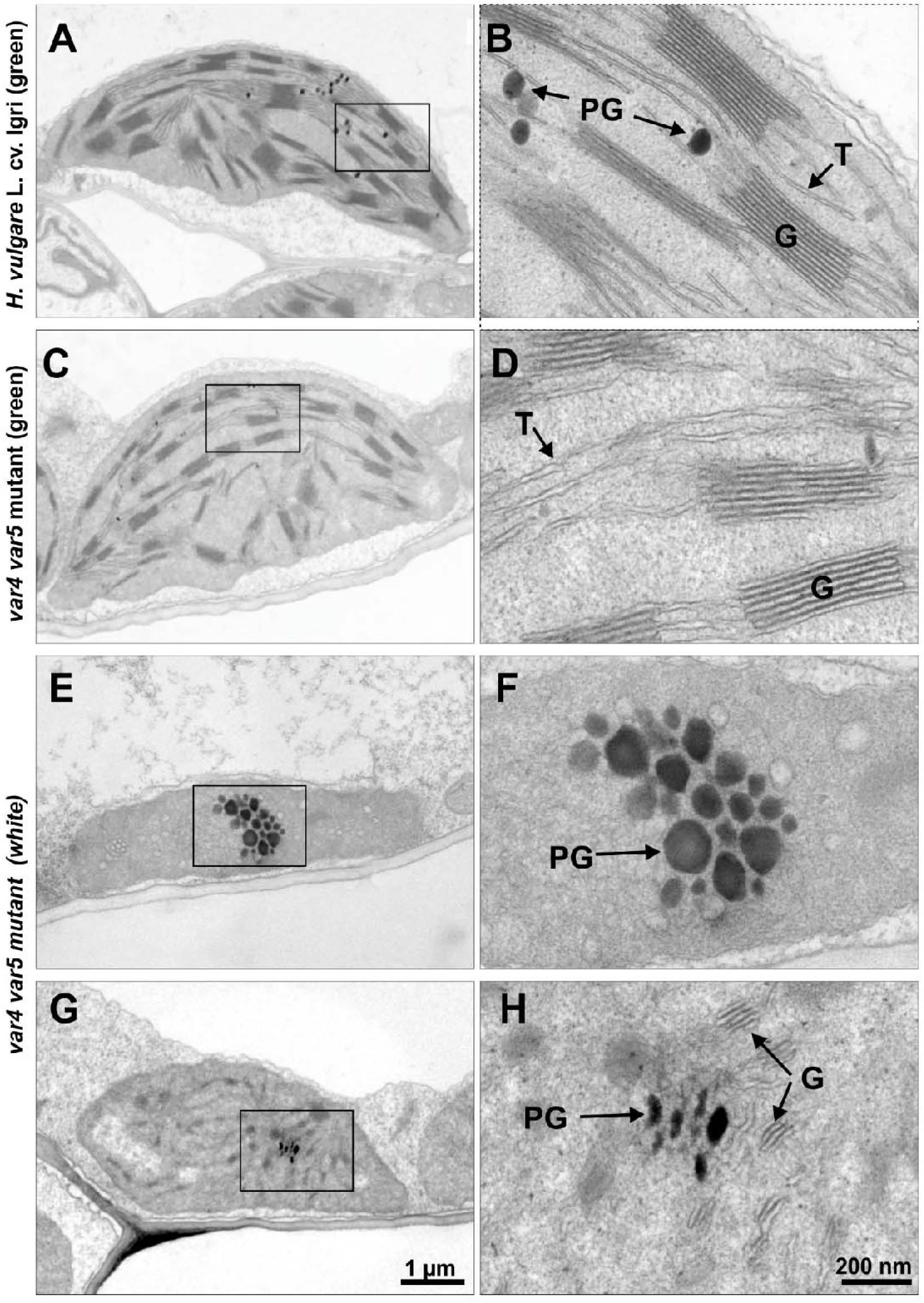
Chloroplast ultrastructure of WT Igri plant and *var4 var5* double mutant 6 months after sowing. Chloroplast with close up of Igri **(A, B)**, green **(C, D),** and yellow sectors of *var4 var5* double mutant **(E, F, G, H)**, respectively. G, grana; ST, stroma thylakoid; PB, prolamellar body; PG, plastoglobuli.

## Discussion

Identification of the genetic basis of variegation is an important tool for a better understanding of chloroplast biogenesis. Here we studied a newly identified variegated barley mutant in which we showed that variegation is controlled by duplicate dominant epistasis, the WT alleles being dominant. To our knowledge, this is the first time that a digenic control has been identified for chloroplast-deficient mutants. However, pairs of genes have been identified as playing a role in variegation, such as *VAR2*/*FtsH8,* which code for proteins part of the same oligomeric complex (Yu *et al*., 2004). The phenotype caused by *var2* is also enhanced by other chloroplastic genes that interact with it (Wang *et al*., 2018; Qi *et al*., 2020). The genetic control of the mutant variegation could be explained by an additive effect of the two mutations combined with a threshold-dependent mechanism similar to the one described for the *Arabidopsis* locus *Var2* (Takechi *et al*., 2000; Yu *et al*., 2007) and the barley mutant *albostrians* (Li *et al*., 2019).

A distinctive feature of the reported mutant here is that the initially white stripes of the early developmental stage turn visibly yellow after about 6 months under greenhouse conditions. Ultrastructural analysis of the chloroplasts in the white sectors at early developmental stages show prolamellar bodies, barely developed thylakoids, and grana. The chloroplast development in those cells appears to be clearly disturbed. After six months, a proportion of the plastids in cells of the yellow sectors have better formed thylakoids and small grana, although these are much less developed than in the green leaf sectors of the mutant or in the WT. These observations suggest a partial recovery of the chloroplast biogenesis process at a later plant developmental stage, possibly caused by the expression of other genes or a shift in the postulated activity threshold level.

The two new genes, *Var4* and *Var5*, were mapped with low resolution on chromosomes 2H and 3H, in 425.3 and 306.6 Mbp intervals, respectively. These intervals do not overlap with the ones described for other variegation loci in barley (Li *et al*., 2019, 2021; Overlander-Chen *et al*., 2024). Using WGS of variegated and green progenies of the mutant backcrossed with the WT, we were able to identify five candidate genes for each of the two intervals on the basis of their sequence variant profile. Interestingly, while functional mutations generated by EMS are usually linked to loss of function, all but one of the candidate mutations are amino acid substitutions. This could account for the unusual detection of two loci in epistasis, as their mild effects may only pass a threshold and produce a visible phenotype when combined.

At *Var4* locus, HORVU.MOREX.PROJ.2HG00154410 emerges as the most likely candidate gene. It is homologous to *A. thaliana* AT4G24690, which encodes NBR1, a selective autophagy receptor and adaptor. NBR1 has been shown to recognize and mediate the degradation of photodamaged chloroplasts via a microautophagy-like process, thereby preventing the accumulation of dysfunctional organelles under high light conditions (Lee *et al*., 2023). In this pathway, the ubiquitin-binding UBA2 domain of NBR1 interacts with the ubiquitinated surface of photodamaged chloroplast and promotes their delivery to the vacuole for degradation. NBR1 also contributes to chloroplast protein homeostasis (proteostasis) by regulating the abundance of TOC components (Wan *et al*., 2023). Changes in TOC protein levels affect the rate of protein import into chloroplasts and consequently, the capacity of these organelles to adapt to different environmental conditions such as heat stress and UV-B radiation. In our barley mutant, alteration of NBR1 function could impair the clearance of damaged chloroplasts—leading to the accumulation of aberrantly developed organelles in specific cell types—or disrupt the adjustment of protein import in response to environmental conditions, thereby compromising chloroplast proteostasis.

At the *Var5* locus, HORVU.MOREX.PROJ.3HG00238870 is the strongest candidate. This gene encodes a DNAJ-domain protein, a member of a co-chaperone family that functions as a specificity factor for Hsp70 chaperones (Pulido and Leister, 2018). DnaJ proteins recognize unfolded or misfolded substrates and deliver them to Hsp70s, which then mediate either their refolding or degradation. In chloroplasts, numerous DnaJ proteins have specialized roles in protein import, folding, assembly, and repair of essential photosynthetic and metabolic complexes. A well-characterized example is AtJ20, a plastid J-protein that specifically binds misfolded deoxyxylulose 5-phosphate synthase (DXS), an aggregation-prone enzyme, and delivers it to chloroplastic HSP70s (cpHSP70s) (Pulido *et al*., 2013). These Hsp70-bound DXS proteins can be routed either to the ClpC1 protease for degradation in a housekeeping pathway or refolded by the disaggregase ClpB3 under stress conditions such as heat (Pulido *et al*., 2016). This DnaJ–HSP70–Clp network forms a central hub of chloroplast proteostasis, preventing the accumulation of toxic protein aggregates and maintaining the functional integrity of plastid enzymes and complexes. Other plastid-localized J-proteins exhibit diverse roles critical for the chloroplast. For example, DJA5 and DJA6 are essential for iron–sulfur cluster biogenesis and plastid morphology (Zhang *et al*., 2021), while AtJ8, AtJ11, and AtJ20 support photosynthetic efficiency by assisting cpHsp70s in the correct folding and assembly of photosystem components and Rubisco (Portis *et al*., 2008; Chen *et al*., 2010). In our barley mutant, disruption of HORVU.MOREX.PROJ.3HG00238870 could compromise the chloroplast’s capacity to maintain protein quality control. This may lead to the accumulation of misfolded or damaged proteins, a collapse in the balance between refolding and degradation, and ultimately a breakdown of chloroplast proteostasis and organellar function in affected cells.

Although these two candidates do not appear to interact directly or participate in the same pathway, both may be involved in chloroplast proteostasis. The chloroplast proteome comprises around 3,000 proteins, of which ∼90% are encoded in the nucleus and must be imported into the chloroplast. Proteostasis is essential to ensure correct protein folding, assembly, and turnover, thereby maintaining the functionality of photosynthetic and metabolic machinery (Jarvis and López-Juez, 2013). While each mutation alone might cause only a mild impairment of proteostasis, their combined effects could disrupt it sufficiently to alter chloroplast morphology and function, which would explain the epistasis we observed. Because the plastid proteome can vary considerably with developmental stage (Teng *et al*., 2012), these effects might be less pronounced in older plants, potentially accounting for the recovery we observed.

While we previously demonstrated that genetic resequencing can be highly effective for identifying strong candidate genes (Li *et al*., 2021; Jhingan *et al*., 2025, Preprint), our analysis focused only on mutations with a direct effect on the protein sequence of these genes. However, the epistatic interaction between two loci complicates the dissection of the genetic control of this variegation, and regulatory regions—excluded from our initial analysis for feasibility reasons—could also contribute. A finer mapping of each locus in F₃ populations, in which the counterpart locus is fixed, would help to confirm or refute our candidates, which in any case will require functional validation.

## Supplementary data

Supplementary Figure 1: Sequence logo of the alignment of the orthogroups of candidate genes for *var4* (A) and *var5* (B) in the 101 bp interval around the polymorphisms carried by the *var4 var5* double mutant.

Supplementary Table S1: PACE markers developed for the fine mapping of *var4* and *var5*mutations.

Supplementary Table S2: Phenotype of the F_2_ plants studied.

Supplementary Table S3 : Genotype and genetic map of the populations AM and MA combined.

Supplementary Table S4: LOD scores of simple (sim) and composite interval mapping (cim). Supplementary Table S5: PACE genotyping of 89 selected F_2_ plants at *Var4* and *Var5* loci.

Supplementary Table S6: Genotype at the *Var4* locus of 30 F2 plants from the IM population mapped on MorexV3 genome.

Supplementary Table S7: Genotype at the *Var4* locus of 30 F2 plants from the IM population mapped on IgriV2 genome.

Supplementary Table S8: Genotype at the *Var5* locus of 30 F2 plants from the IM population mapped on MorexV3 genome.

Supplementary Table S9: Genotype at the *Var5* locus of 30 F2 plants from the IM population mapped on IgriV2 genome.

## Acknowledgements

We gratefully acknowledge excellent technical support from Mary Ziems for the population construction and greenhouse management; Mary Ziems and Pascal Jaroschinsky for plant phenotyping and dna extraction; Ines Walde, Jaqueline Pohl, Marius Dölling and Susanne König for library preparation and Illumina sequencing; Marion Benecke, Claudia Riemey and Kirsten Hoffie for technical assistance with light and electron microscopy. We thank Anne Fiebig for data submission to ENA and Mats Hansson for his input on the naming of the two mutations.

## Author contributions

NS: conceptualization; HP: data curation; HP, SRN, and MM: formal analysis; HP, SRN, MM, and AH: investigation; HP, and NS: writing – original draft preparation; HP, NS, SRN, MM, and AH: writing – review & editing; NS: project administration; NS: funding acquisition

## Data availability

All raw sequencing data produced for this paper is available under projects number PRJEB104329 (GBS data), and PRJEB104329 (WGS data).

## Abbreviations

AM: Alraune x mutant
GBS: Genotyping-by-sequencing IM: Igri x mutant
MA: mutant x Alraune
PACE: PCR Allele Competitive Extension
TEM: transmission electron microscopy
*Var4*: Variegated 4
*Var5*: Variegated 5
WGS: whole-genome shotgun sequencing

